# Inactivation effects of plasma-activated water on *Fusarium graminearum*

**DOI:** 10.1101/2021.07.15.452455

**Authors:** Jian Guo, Jiaoyu Wang, Hui Xie, Junlong Jiang, Chunyuan Li, Wanting Li, Ling Li, Xingquan Liu, Fucheng Lin

## Abstract

The continuous usage of fungicides poses a potential threat to the environment, ranging from mere irritation to being very toxic to human beings and organisms. Plasma-activated water (PAW) has recently gained much interest as a promising candidate to inactivate fungi. However, the inactivation mechanisms of PAW are still not well understood. In this study, the effect of PAW on the viability and the cellular responses of *Fusarium graminearum* in PAW inactivation were investigated. The results showed that microbial activity of spores was significantly inhibited by PAW treatment (*P* < 0.05). The symptoms caused by *F. graminearum* were significantly reduced on the spikelets. Our data indicated that PAW could induce cell wall sculpturing, membrane permeability changes, and mitochondrial dysfunction. Differential gene expression analysis also confirmed that the cell membrane, the cell wall and the mitochondria were the organelles most affected by PAW. The results from this study facilitate the understanding of the mechanisms underlying the responses of *F. graminearum* to PAW and the development of PAW as a potential fungicidal agent or an effective supplement to fungicides.

**Highlights:**

- The viability of F. graminearum is notably inhibited by PAW
- The symptoms caused by *F. graminearum* were significantly reduced on the spikelets
- Oxidative stress induce cell wall sculpturing, membrane permeability change
- PAW can cause the mitochondrial dysfunction
- Cell wall, membrane and mitochondria are the most affected organelles by PAW

## 1. Introduction

Phytopathogens are the main causes of plant diseases, they significantly reduce the yield of agricultural crops (McMullen et al., 1997; Marcos et al., 2008). The use of fungicides in modern agricultural practices has made crop production stable, and environmental and food safety issues associated with the use of fungicides cannot be ignored. Prolonged or repeated application of agricultural chemicals has led to the outbreak of acute or chronic diseases, toxicity to nontarget organisms, excessive fungicide residues, and groundwater and surface water contamination (Pimentel et al., 1992; Soylu et al., 2010). *Fusarium graminearum* is an ascomycetous fungus that is the major cause of *Fusarium* head blight (FHB) on wheat and barley, as well as stalk and ear rot disease on maize. This fungus not only causes crop yield and quality losses but also contaminates grain with mycotoxins, rendering them unfit for food or feed (Etzerodt et al., 2016; Kim and Vujanovic, 2016; Masci et al., 2015; Zhou et al., 2021). Benzimidazoles and sterol demethylation inhibitors are widely used for the control of FHB (Blandino et al., 2006; Bian et al., 2021). However, these synthetic chemicals are difficult to degrade and are harmful to the environment and human health (Zheng et al., 2019). Therefore, much attention has been given to the development of highly efficient, environmentally friendly, and low-residue novel antifungal agents. Cold plasma is an electrically energized matter in a gaseous state that is composed of charged particles, free radicals, and some radiation. To obtain cold plasma, an electrical discharge device is needed, which is not convenient for practical applications. Plasma treatment of water, termed plasma-activated water (PAW), alters the physicochemical properties of water, such as the redox potential, pH and conductivity. As a result, PAW has a different chemical composition than water and can serve as an alternative method for microbial disinfection (Ercan et al., 2013; Guo et al., 2021; Natali et al., 2010; Oehmigen et al., 2010; Puligundla et al., 2018; Shen et al., 2016; Traylor et al., 2011; Tian et al., 2015; Xiang et al., 2019a; Xu et al., 2020; Zhang et al., 2016). Reactive oxygen and nitrogen species (RONS) are the major agents for air-plasma-induced biological effects, and almost no harmful byproducts are generated in the plasma treatment process (Xu et al., 2020). In our previous study, the oxidative stress and acidity of PAW were not persistent (Guo et al., 2017; Guo et al., 2021; Shen et al., 2016). The lifetime of the chemical components in PAW is short and labile (Brisset et al., 2011; Girard et al., 2016; Pavlovich et al., 2013), and it leaves less residue and harmful chemical components on crops than conventional approaches. Thus, PAW can be regarded as a promising eco-friendly approach for fungal inactivation.

Previous studies have showen that chemical components generated in PAW and the oxidative stress it induces play major roles during PAW treatments (Natali et al., 2010; Oehmigen et al., 2010; Pavlovich et al., 2013; Zhang et al., 2013). It is generally accepted that different plasma devices and working gases could lead to different inactivation efficiencies and inactivation patterns. For instance, oxygen-containing gases can increase the proportion of reactive oxygen species (ROS) leading to a higher inactivation efficiency. It was also found that ROS play an important role in direct plasma inactivation in an aqueous environment (Chandana et al., 2018; Gorbanev et al., 2018; Guo et al., 2021; Xu et al., 2020). Kaneko et al. (2017) found that ·OH plays a key role in gas-liquid interfacial plasma irradiation. Xu et al., (2020) reported that ^1^O_2_ contributes the most to the yeast cells. Because of the complexity of the reactive species in plasma and the different roles played by short- and long-lived ROS, the cold plasma inactivation mechanisms caused by these RONS in an aqueous environment or PAW are not yet fully understood.

Apart from the different roles played by the diverse reactive species in the cold plasma inactivation process, the complexity of the antimicrobial mechanisms of PAW is also attributed to the complicated cellular response. Many works have found that different types of microorganisms have different responses to cold plasma treatment or PAW treatment. Many works have reported that different types of microorganisms behave differently in their interactions with RONS in cold plasma or PAW. Xiang et al. (2019b) found that gram-negative *E. coli* bacteria were more sensitive to PAW than the gram-positive *S. aureus* bacteria, which might be due to the significant differences in the cell wall structures of gram-negative and gram-positive bacteria, especially the thickness of the peptidoglycan layer. Han et al. 2016 reported that cold plasma inactivated *E. coli* mainly through damaging the cell wall, while *S. aureus* was inactivated by cold plasma primarily through damaging the intracellular components. Furthermore, it is commonly considered that fungal resistance against cold plasma is generally higher than that in bacteria due to the complex cell wall structure and specialized cellular organelles (such as mitochondria and ribosomes) (Xu et al., 2020). The cell wall of fungi consists of chitin, which is more rigid than the peptidoglycan of bacterial cell walls. Lunov et al. (2016) reported that plasma could induce two different inactivation mechanisms (apoptosis or direct physical destruction) in bacteria depending on the plasma treatment time. Xu et al. (2020) also revealed that yeast cells underwent apoptosis in the first 3 min of treatment due to the accumulation of intracellular ROS, mitochondrial dysfunction and intracellular acidification, followed by necrosis under longer exposure times, attributed to the destruction of the cell membrane. Itooka et al. (2018) revealed that cold plasma could cause protein denaturation and endoplasmic reticulum stress in *Saccharomyces cerevisiae*. However, studies that focus on the cellular responses in the PAW inactivation of fungi are very limited, especially from the perspective of gene expression.

In this work, the efficacy of PAW against *F. graminearum in vivo* and *in vitro* was first estimated by calculating the disease severity index and assessing the effects of PAW on mycelium growth, conidium germination and conidiation. The complicated cellular responses of *F. graminearum* to PAW were explored by assessing cell morphology, cell membrane integrity, mitochondrial activity and gene expression.

## 2. Materials and methods

### 2.1. Fungal culture conditions

The *Fusarium graminearum* PH-1 strain used in this study was obtained from the Agricultural Culture Collection of China (Beijing, China). *F. graminearum* was aseptically inoculate in liquid complete medium (CM) at 30 °C for 7 d. The strain was routinely cultured on CM plates at 28 °C for mycelial growth assays. For the conidial germination assay, three mycelial plugs of the strain were inoculated in 30 mL of liquid CM under continuous light. After incubation in a shaker at 180 rpm at 28 °C for 7 d, conidia of each sample were collected by centrifugation and were quantified using a hemocytometer.

### 2.2. Plasma treatment system

The cold plasma device and operation method were described in our previous work (Guo et al., 2021). Sterile distilled water (200 mL) was activated by plasma treatment for 15, 30, 45 and 60 min. Accordingly, the samples after treatment were named as PAW15, PAW30, PAW45 and PAW60, and the control.

### 2.3. Physicochemical properties of PAW

To evaluate the physicochemical properties of PAW, the conductivity, oxidation reduction potential (ORP), and pH of PAW were all detected using a multimeter (Orion 3 Star pH/ORP Meter, Thermo Fisher Scientific Inc, PA) immediately after production. ORP was used to estimate the oxidative stress in PAW, the concentration of the oxidizers and their activity or strength (Mcferson, 1993).

The concentrations of nitrate anions (NO_3_) and nitrite anions (NO_2_) in PAW were determined by spectrophotometry (Shen et al., 2016; Collos et al., 1999). Then, 50 mL of PAW was added to 1 mL of 1 M hydrochloric acid and 100 μl of 0.8% sulfamic acid. NO_3_ levels in PAW samples were determined by ultraviolet absorption spectrometry (UV-1800, Shimadzu Corporation, Kyoto, Japan) at a single wavelength of 220 nm.

For the determination of NO_2_, sulphanilamide was used as the diazotizing reagent, and N-(1-naphthy1)-ethylenediamine hydrochloride was used as the coupling reagent. After plasma activation, 1 mL of sulphanilamide solution (5 g of sulphanilamide was dissolved in a mixture of 50 mL of 37% (w/w) concentrated HCl and 300 mL of water, and it was diluted to 500 mL with water) was added into 50 mL of PAW, and incubated at room temperature for 2 min, subsequently adding 1 mL of 1.0 g L^-1^ N-(1-naphthy1)-ethylenediamine hydrochloride at room temperature. After a 20-min incubation at room temperature, the absorbance at 540 nm was measured using a spectrophotometer (UV-1800, Shimadzu Corporation, Kyoto, Japan) for the determination of the concentrations of NO_3_ in the PAW.

### 2.4. In vivo fungicidal activities assay

The *in vivo* fungicidal activities of PAW against *F. graminearum* (strain PH-1) were evaluated as follows: The spikelets were point-inoculated with 20 μl of the conidial suspension (10^5^ conidia mL^-1^) mixed with PAW. After inoculation, the spikelets were placed in a controlled chamber (90‒100% relative humidity, 22 °C) for 72 h. The disease severity was scored by measuring the lesion area with image analysis software (ImageJ), and the disease severity index was calculated (the mean area of disease divided by the total area of the wheat × 100%). The protection efficacies were calculated as follows: Protection efficacy = [(mean area of infected spikelets of control – mean area of infected spikelets of treated group)/mean area of infected spikelets of control] × 100%.

### 2.5. In vitro fungicidal activities assay

For the mycelium growth assay, 2 mL of PAW was transferred into the test tubes containing 0.5 mL of *F. graminearum* spore suspension. After exposure for 1 h, 10 μl of *F. graminearum* spore suspension was dropped at the center of CM or CR (Congo red was added to complete medium to 100 µg mL^-1^ or 200 µg mL^-1^) and CFW (calcofluor white was added to complete medium to 50 µg mL^-1^) plates. The agar plates were incubated at 28 °C. Sterile water (2 mL) was used as the control. The colonial diameters of the treated strain were carefully measured by a caliper, every 12 h over 5 d, and the inhibition ratios were calculated using the following formulas:

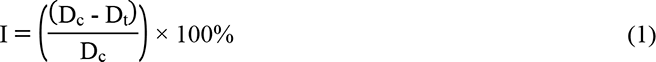

where D_c_ and D_t_ are the colonial diameters of the control and treated groups, respectively.

The dry weight of the mycelial biomass was measured after 7 days of cultivation in liquid CR or CFW medium.

Conidiation was assessed by growing the strain in liquid complete medium (CM) for 7 days. The conidial concentration was measured using a hemocytometer.

For the conidial germination assay, spores from strain PH-1 were inoculated in the liquid CM in the presence of PAW or were left untreated. PAW15, PAW30, PAW45 and PAW60 (0.4 mL) were transferred into tubes that contained an *F. graminearum* spore suspension (0.1 mL), and the mixture was allowed to stand for 1 h. Five microliters of the treated conidia was pipetted onto the concave slide at 25°C for 6 h and then observed by fluorescence microscopy (Nikon Eclipse Ti-s; Tokyo, Japan) for the analysis of germ tube emergence. Sterile water (0.4 mL) was used as the control. Three different fields of view were observed randomly. The number of conidia observed in each test was at least 200. Although germ tube emergence marks the culmination of the germination process, it was used as a convenient marker for germination in our experiments (Semighini et al., 2008). The conidial germination ratios were calculated using the following formulas:

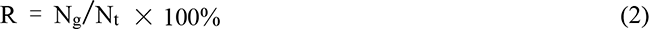

where R is the conidial germination ratio, N_g_ is the conidial germination number, and N_t_ is the conidial number.

The relative conidial germination ratios were calculated using the following formula:

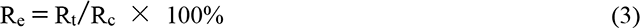

where R_e_ is the relative conidial germination ratio, and R_t_ and R_c_ are the conidial germination ratios of the treated and control groups, respectively.

### 2.6. Effect of PAW on cell wall integrity

The effect of PAW on cell wall integrity was assayed by measuring the conidiation and dry mycelial weight of the strain grown in liquid CM supplemented with 100 mg L^-1^, 200 mg L^-1^ Congo red and 50 mg L^-1^ Calcofluor White for 7 days.

### 2.7. SEM and TEM observations

To investigate the effect of PAW on the spore cell wall. We performed transmission electron microscopy (TEM) imaging. After PAW treatment, the spores were fixed in Karnovsky’s fixative (2% v/v paraformaldehyde and 2% v/v glutaraldehyde in 1x PBS) overnight. The specimens were observed using a Hitachi Model H-7650 TEM (Hitachi High-Tech Manufacturing & Service, Corp).

### 2.8. Measurement of nucleic acid and protein leakage

Spore cell membrane integrity was detected by determining the release of intracellular components absorbing at 260 nm and 280 nm as described in the relevant literature (Cockrell et al., 2017; Xiang et al., 2018) with slight changes. Spores were harvested by centrifugation at 5000 x g for 10 min at 4 °C, and were then suspended in sterile deionized water. PAW15, PAW30, PAW45 and PAW60 were added to spores suspensions. Spores treated with PAW0 were used as the control. To quantify the concentrations of nucleic acids and proteins, the optical densities of the supernatants at 260 nm and 280 nm were recorded.

### 2.9. Measurement of mitochondrial activity and total microbial activity

Spores were labeled with tetramethylrhodamine methyl ester (TMRM, 80 nM) for mitochondria after PAW treatment. The cells were then washed three times and resuspended in PBS for observation. Imaging of cells loaded with fluorescent dyes was performed using a Zeiss LSM510 system (Carl Zeiss, Germany). The excitation wavelength was 543 nm and the emission was collected between 561 and 603 nm.

A stock solution of 500 μg mL^-1^ FDA was made in acetone and then diluted to 100 μg mL^-1^ in incubation buffer. The cells were then washed three times and resuspended in PBS for observation. Fluorescein was excited using a 488 nm laser, and the emission was collected between 505 and 530 nm.

### 2.10. RNAseq and transcriptomic analyses

Fungal materials were frozen in liquid nitrogen and RNA was extracted using TRIzol^®^ Reagent according to the manufacturer’s instructions (Invitrogen), and genomic DNA was removed using DNase I (TaKaRa). The RNA-seq transcriptome library was prepared following the TruSeq^TM^ RNA sample preparation kit from Illumina (San Diego, CA) using 1 μg of total RNA. After quantification by TBS380, the paired-end RNA-seq sequencing library was sequenced using an Illumina HiSeq xten/NovaSeq 6000 sequencer (2 × 150 bp read length). To identify DEGs (differentially expressed genes) between two different samples, the expression level of each transcript was calculated according to the fragments per kilobase of exon per million mapped reads (FRKM) method. RSEM was used to quantify gene abundances (Li and Dewey 2011). Differential gene expression analysis was performed with DESeq2 software based on negative binomial distribution.

### 2.11. Quantitative PCR

To evaluate the validation of RNA-seq transcriptome results, primers (Table 2) were designed for the following thirteen candidates selected based on the following criteria: the highest upregulated gene involved in mitochondrial function and the highest upregulated or the lowest downregulated gene involved in cell wall and membrane integrity. Quantitative PCR (Q-PCR) assays were conducted according to the relevant literature (Liu et al., 2010; Demissie et al., 2018) with slight changes. Normalized relative expression values (ΔΔC_T_) of the selected candidates were calculated using the formula 2^-ΔΔCT^ (Livak and Schmittgen, 2001) using actin as a reference gene. Three amplifications were performed for each replicate. The expression of each tested gene in the PAW-treated sample relative to that of the untreated sample was calculated according to the fragments per kilobase of exon per million mapped reads (FRKM) method.

### 2.12. Statistical analysis

Data from all experiments are expressed as the mean ± standard deviation (SD). At least three replicates were performed for each treatment condition. Statistical analysis was performed using Origin 2019b program. Analysis of variance (ANOVA) was used to calculate significant differences and to compare the means.

## 3. Results and discussion

### 3.1. Physicochemical properties of PAW

It is generally agreed that nitrate and nitrite usually exist in PAW generated by the air-cold plasma devices (Patangea et al., 2019; Pavlovich et al., 2013; Sarangapani et al., 2017). As shown in Fig. 1a, the concentrations of nitrate and nitrite in PAW increased significantly (*P* < 0.05) over air-plasma-activated time. Patangea et al. (2019) also reported that PAW generated using this submerged DBD air-plasma device contained high concentrations of nitrate but a very low concentration of nitrite. When the discharge area is above the liquid surface, similar amounts of nitrate and nitrite were generated in the PAW using this type of cold plasma device (Pavlovich et al., 2013; Sarangapani et al., 2017). When the discharge area is below the water surface, there are more changes for the oxygen or ROS in air plasma to oxidize the nitrite in PAW. Therefore, most nitrite in the PAW generated using a submerged DBD air-plasma device is transformed into nitrate (Guo et al., 2021).

**Fig. 1.**
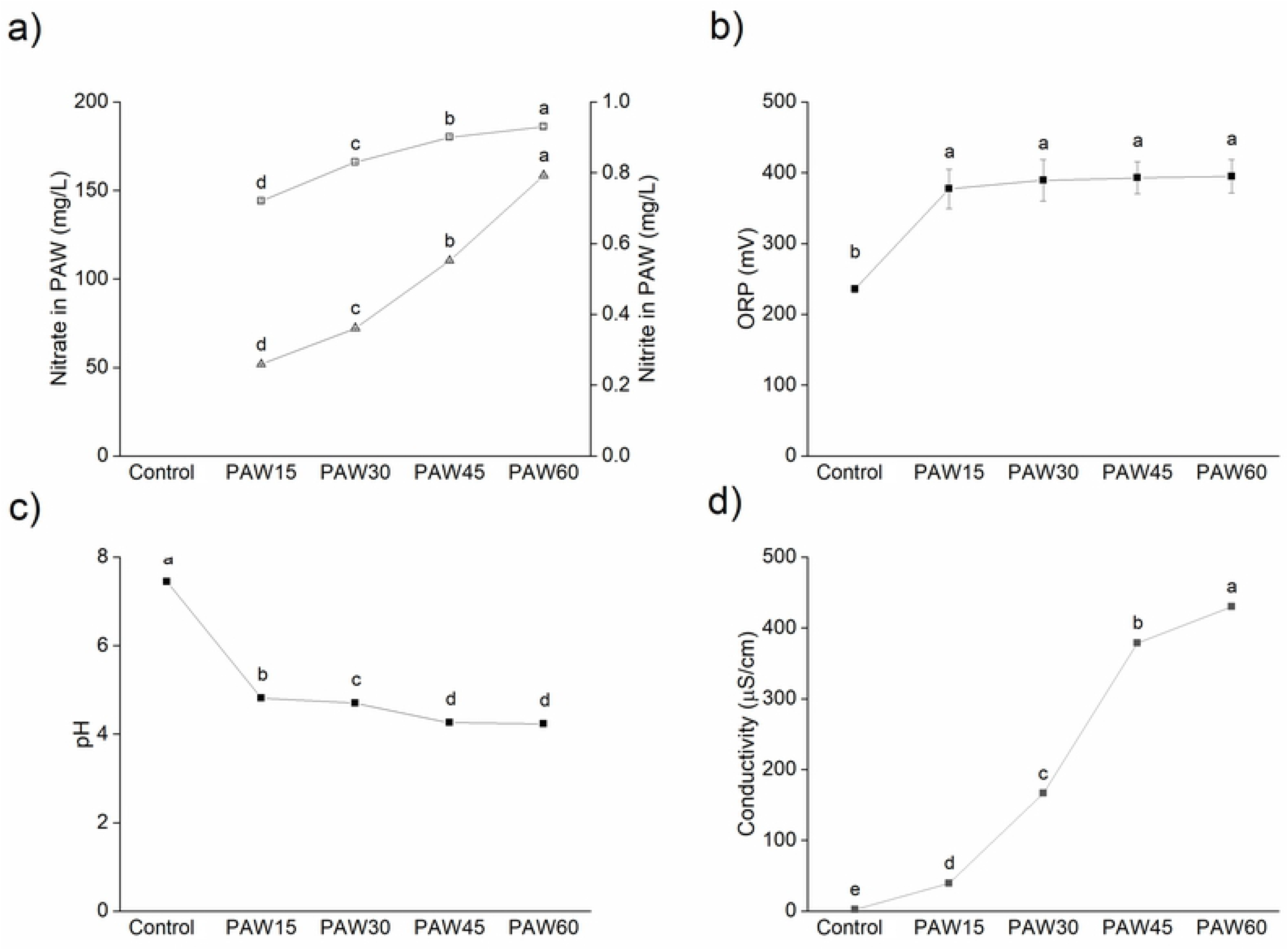
Physicochemical properties of PAW: a) Concentrations of nitrate anions (NO_3_^-^) (□) and nitrite anions (NO_2_^-^)(Δ) in PAW. b) ORP value of PAW subjected to plasma activation for 0, 15, 30, 445, and 60 min. c) pH value of PAW subjected to plasma activation for 0, 15, 30, 45, and 60 min. d) Conductivity of PAW subjected to plasma activation for 0, 15, 30, 45, and 60 min. The results represent the mean±standard deviation (n=3). Vertical bars represent standard deviation of the mean, columns with different letters represent statistically significant results (*P* < 0.05).

ORP reveals the total oxidation ability of various substances in a solution, and is an important indicator in water disinfection applications because there is a direct correlation between high ORP and cell membrane damage (Ma et al., 2015; Tian et al., 2015; Xiang et al., 2018). As shown in Fig. 1b, the ORP of water after plasma activation increased from 235.8±5.43 mV to 394.9±23.69 mV, and an increase in the ORP was observed with prolonged plasma-activation time (*P* < 0.05). Similar results were also obtained in previous work (Guo et al., 2021; Tian et al., 2015; Xu et al., 2016; Xu et al., 2020). The discharged gases in these previous works all contain oxygen, which is the same as our study. Hence, a longer duration of plasma activation results in a larger amount of reactive oxygen species generated in PAW and a high ORP value.

The pH values of PAW over the plasma-activated time were also measured (Fig. 1c). The pH value of water after plasma activation decreased significantly (*P*<0.05) from 7.44±0.010 to 4.234±0.010. However, after 45 min of plasma activation, the extended treatment time did not significantly influence the pH values of PAW, which decreased over the plasma-activated time. Plasma discharge acidified activated water (Oehmigen et al., 2010; Tian et al., 2015; Zhang et al., 2013; Xu et al., 2016). Due to the discharge gases, some nitrate and nitrite were generated in PAW. Prolonged plasma activation corresponds to larger amounts of hydrogen nitrate and nitrous acid, which lead to a lower pH value.

The electrical conductivity can indicate the concentrations of free ions present in an electrolytic solution. The electrical conductivity of PAW was measured before and after plasma activation (Fig. 1d). The conductivity of water after plasma activation increased from 2.3±0.3 μS cm^-1^ to 430.2±1.1 μS cm^-1^. The electrical conductivity of PAW increased in a plasma activation time-dependent manner, which provided evidence for the accumulation of active ions in PAW. The results demonstrated that many active ions were generated in PAW, which may be nitrate acid derived from chemical reactions between electrons or RNS in cold plasma and water molecules (Guo et al., 2021; Ma et al., 2015).

However, the changes in ORP, pH and conductivity were not constant, which may be ascribed to the decomposition of ozone, reactive oxygen species and reactive nitrogen species (Guo et al., 2017; Guo et al., 2021; Shen et al., 2016).

### 3.2. In vivo fungicidal activities assay

To determine the fungicidal activities of PAW against FHB, the protection efficacies and disease severity indices were calculated. The symptoms caused by the pathogen were significantly reduced on the spikelets (Fig. 2). The pathogen treated by PAW generally failed to colonize the inoculated spikelets. The protection efficacies of PAW15, PAW30, PAW45 and PAW60 were 67.8%, 57.4%, 92.7% and 86.8%, respectively. Evaluations of disease development, which were quantified by calculating the disease severity index and the protection efficacy, revealed that PAW exhibited significant fungicidal activity (*P*<0.05) against FHB compared with the control.

**Fig. 2.**
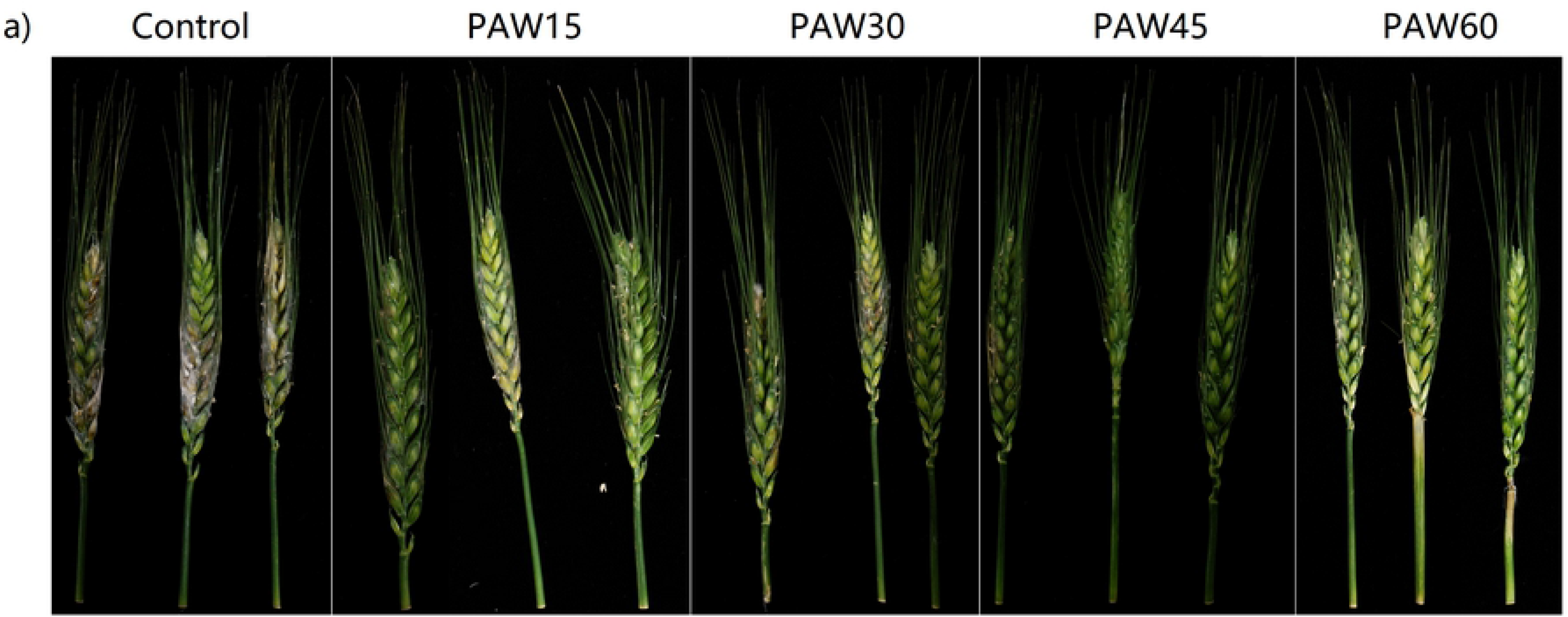
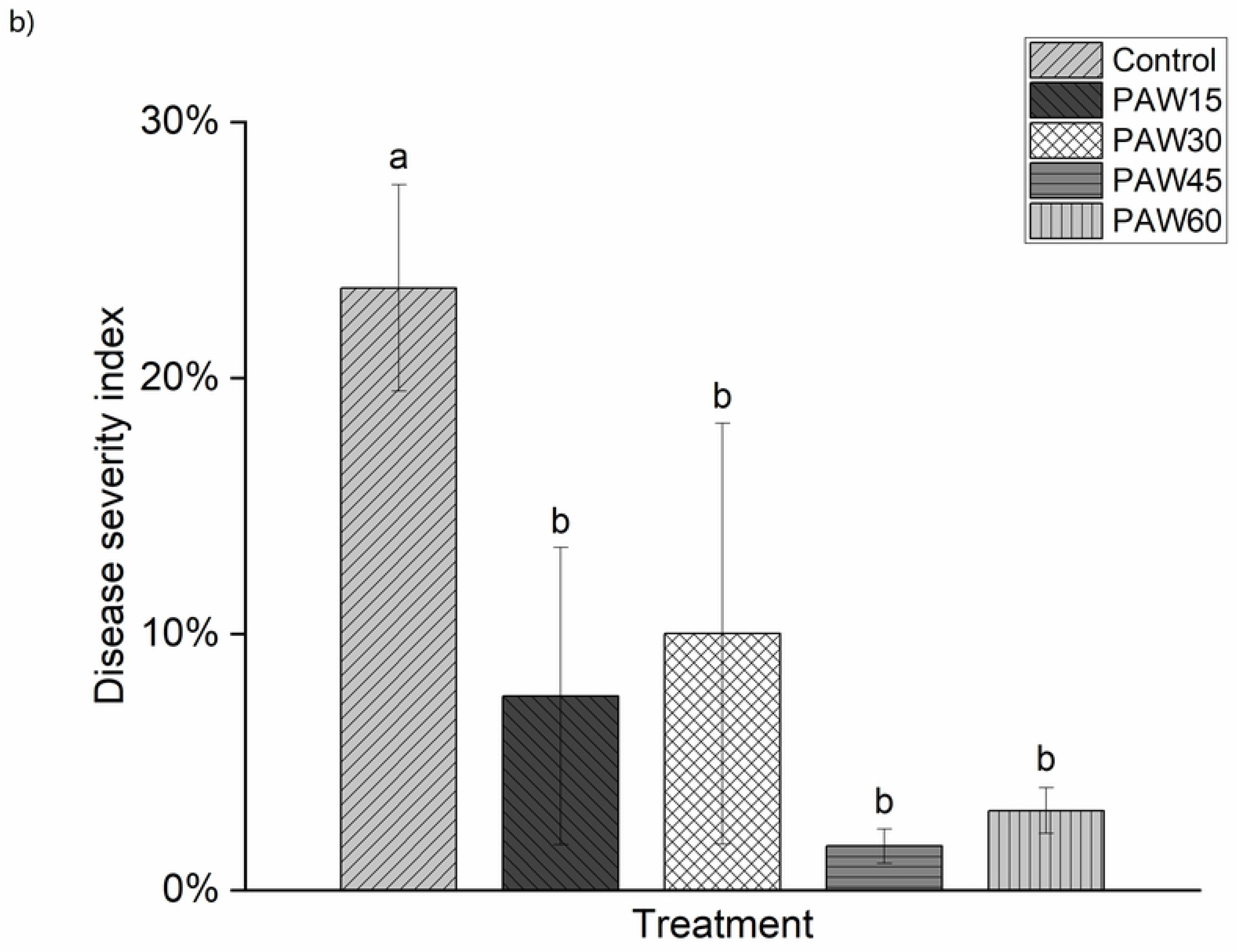
Disease severity indexes are the mean area of disease divided by the total area of the wheat × 100%. Groups of bars followed by the same letter are not significantly different according to Fisher’s protected least significant difference (*P* < 0.05).

The fungicides phenamacril, carbendazim and demethylation inhibitors are commonly used in the field. Compared to chemical synthetic fungicides, PAW has a similar protection efficacy (Chen et al., 2020; D’Angelo et al., 2014). The control efficacy of PAW in FHB increased with increasing the plasma activation time. In a previous study, there was a significant positive correlation between the ORP value and fungicidal activities. The half-value period of ORP in PAW stored inside was estimated to be approximately 9 days (Guo et al., 2021). The half-value period may decrease when PAW is used outdoors. Light can induce the acceleration of nitric acid dissipation of in PAW. The half-lives of phenamacril and carbendazim in soil were found to be 17.1 days and more than 28 days respectively (Donau et al., 2019; Huang et al., 2020). The half-lives of propiconazole and epoxiconazole varied between 20 and 130 days depending on the soil type (Hollomon, 2017). Therefore, in general, PAW has a shorter effective period of antifungal activity than chemical synthetic fungicides.

### 3.3. Effect of PAW on mycelium growth, conidiation, conidium germination and total microbial activity

The results showed that the radial growth of *F. graminearum* was severely (*P* < 0.05) inhibited after PAW treatment (Fig. 3a). After 12 h of incubation, the mycelium length of PAW15-treated *F. graminearum* was 0.33±0.58 mm, whereas that of the control group was 1.56±0.43 mm. The mycelium length of *F. graminearum* did not increase after treatment with PAW30, PAW45, and PAW60. After incubation for 24 h, the radical growth of *F. graminearum* treated with PAW60 was still severely inhibited, but the mycelium lengths of the PAW45, PAW30, and PAW15 groups reached 1.46±0.44, 1.31±0.13, and 3.88±1.16 mm, respectively. After incubation for 36 h, a general increase in radial growth of *F. graminearum* treated with PAW60 was observed. Treatment with PAW led to an approximately 30% reduction in biomass production in comparison to untreated samples (Fig. 3b). The inhibition ratios of mycelium growth are shown in Fig. 3c. Twelve hours after treatment, the PAW30, PAW45, and PAW60 still demonstrated complete inhibition of mycelial growth. Twenty-four hours after treatment, only PAW60 completely inhibited mycelial growth. After 36 h of incubation, the inhibition ratio of mycelium growth in the PAW60 group decreased. These results were in line with those of previous studies showing that fungi were effectively inactivated as the plasma activation time increased (Ercan et al., 2013; Tian et al., 2015; Zhang et al., 2013; Ma et al., 2015). The results suggest that PAW treatment inhibited mycelial growth in a plasma-activated time-dependent manner.

**Fig. 3.**
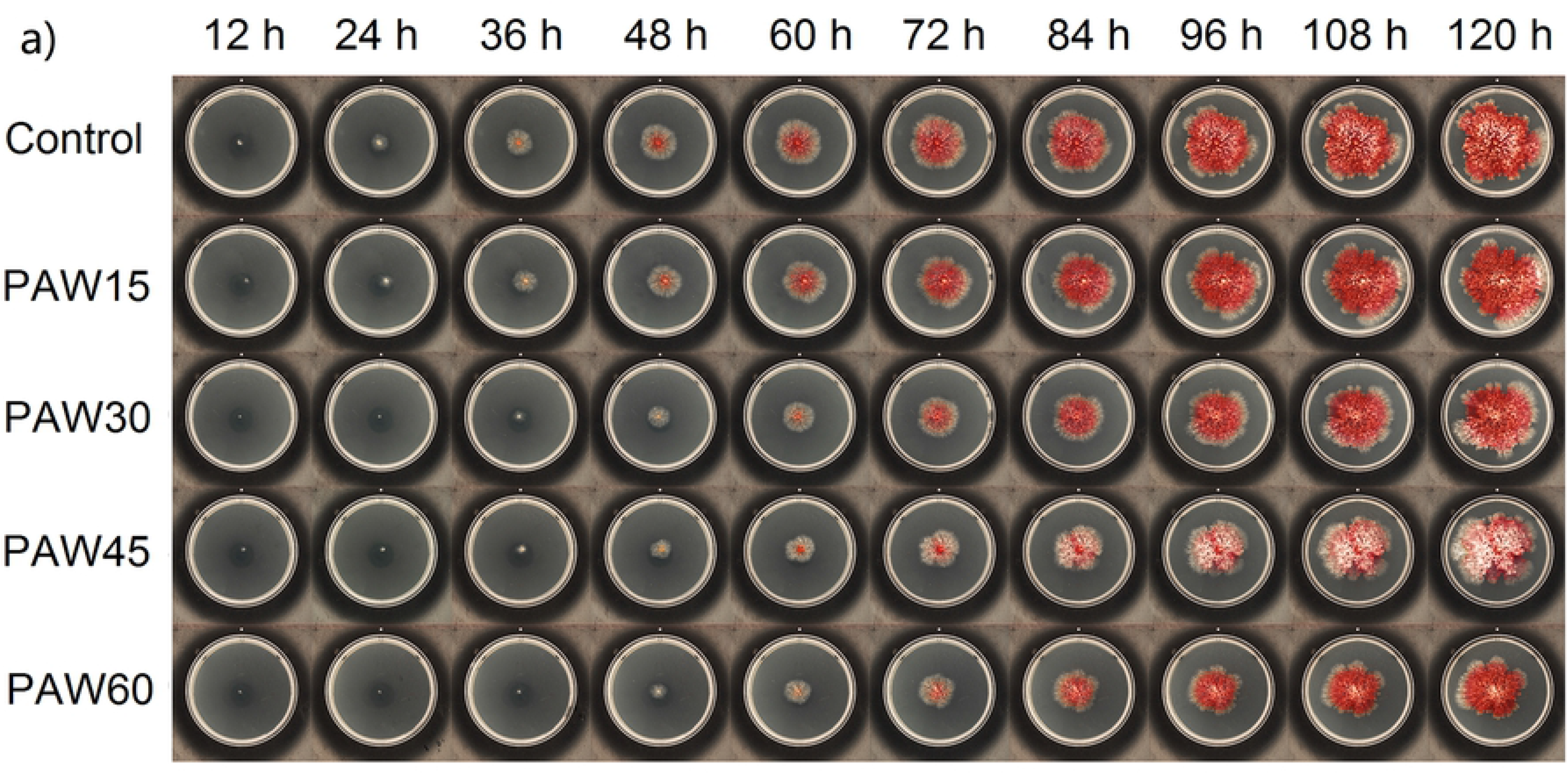
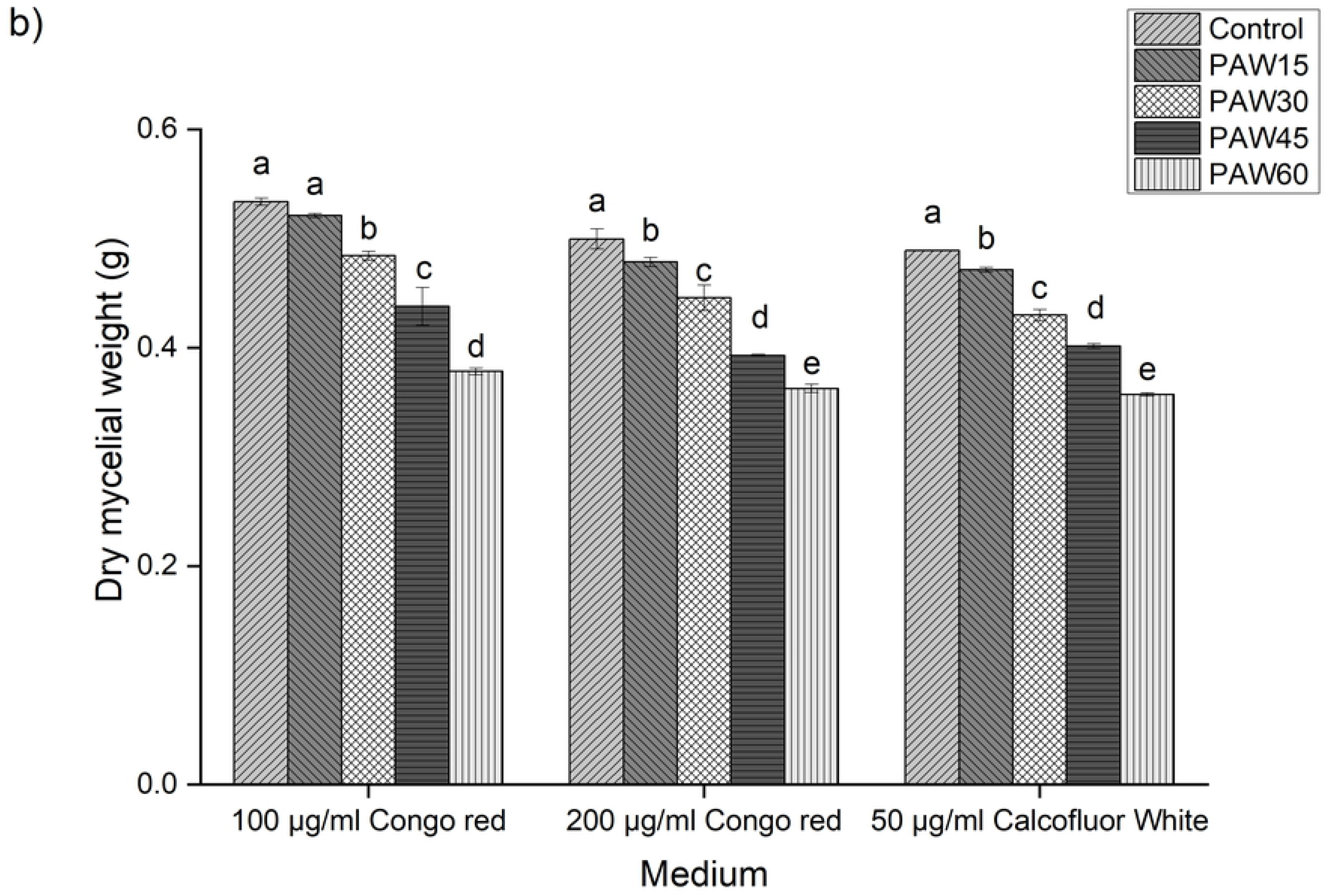
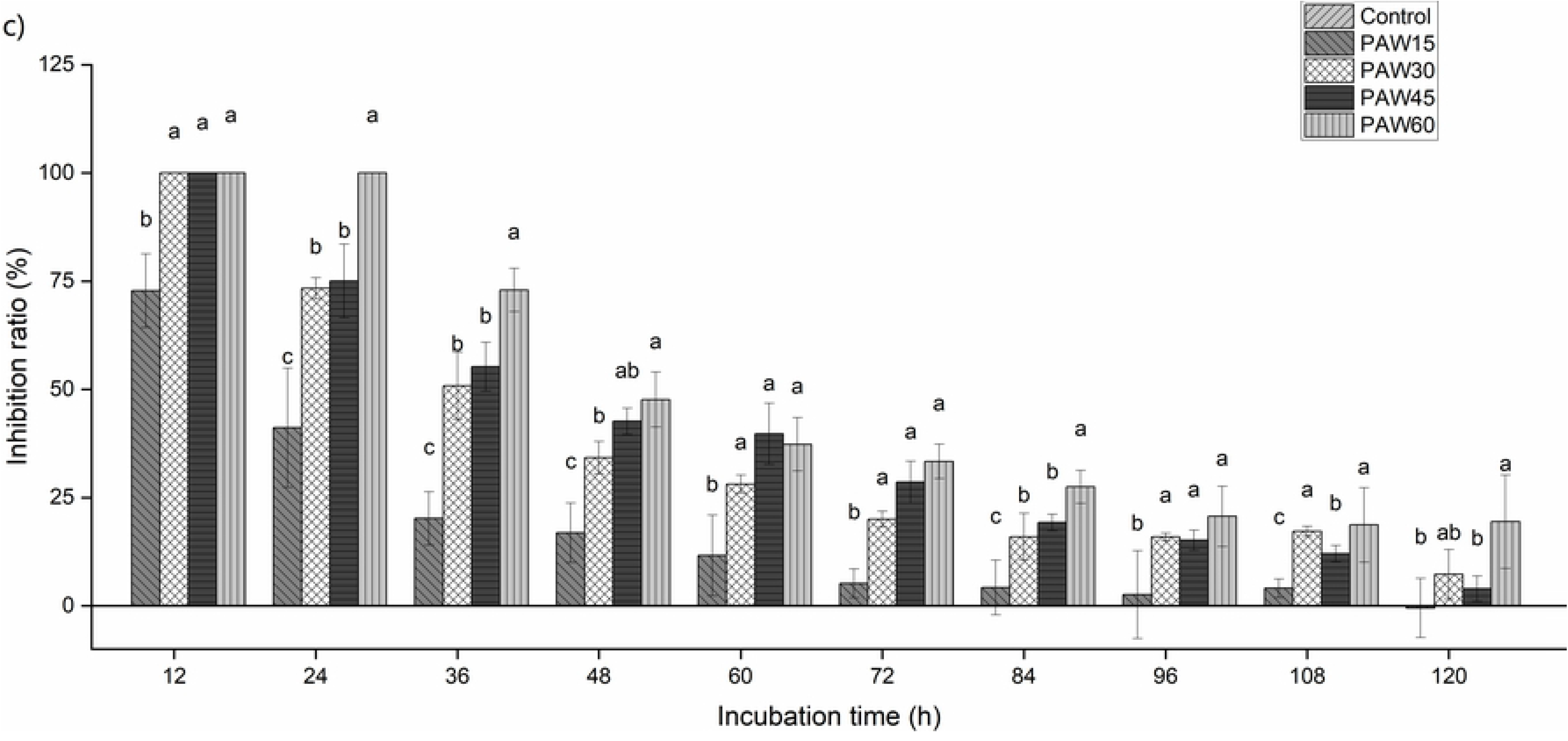
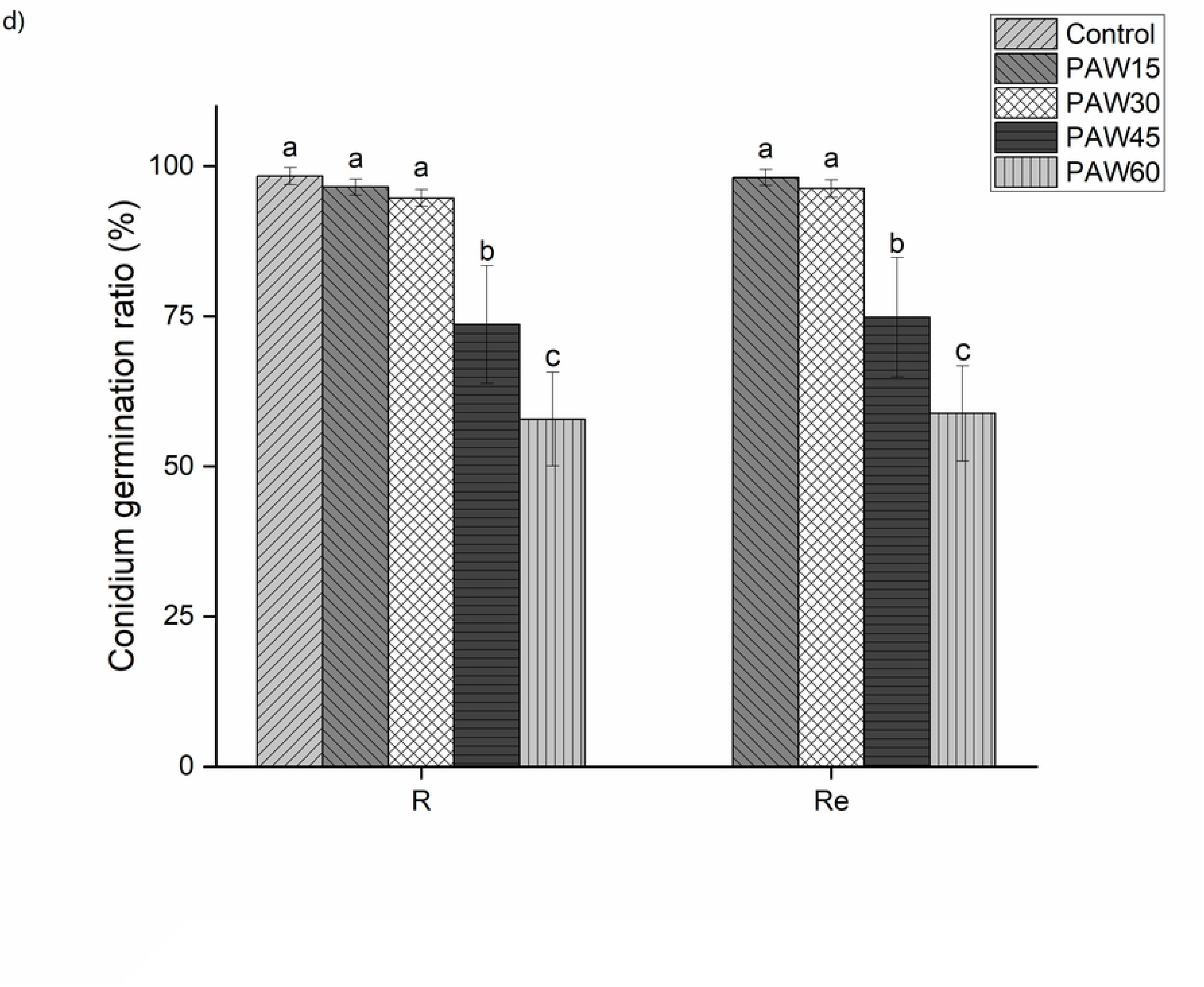
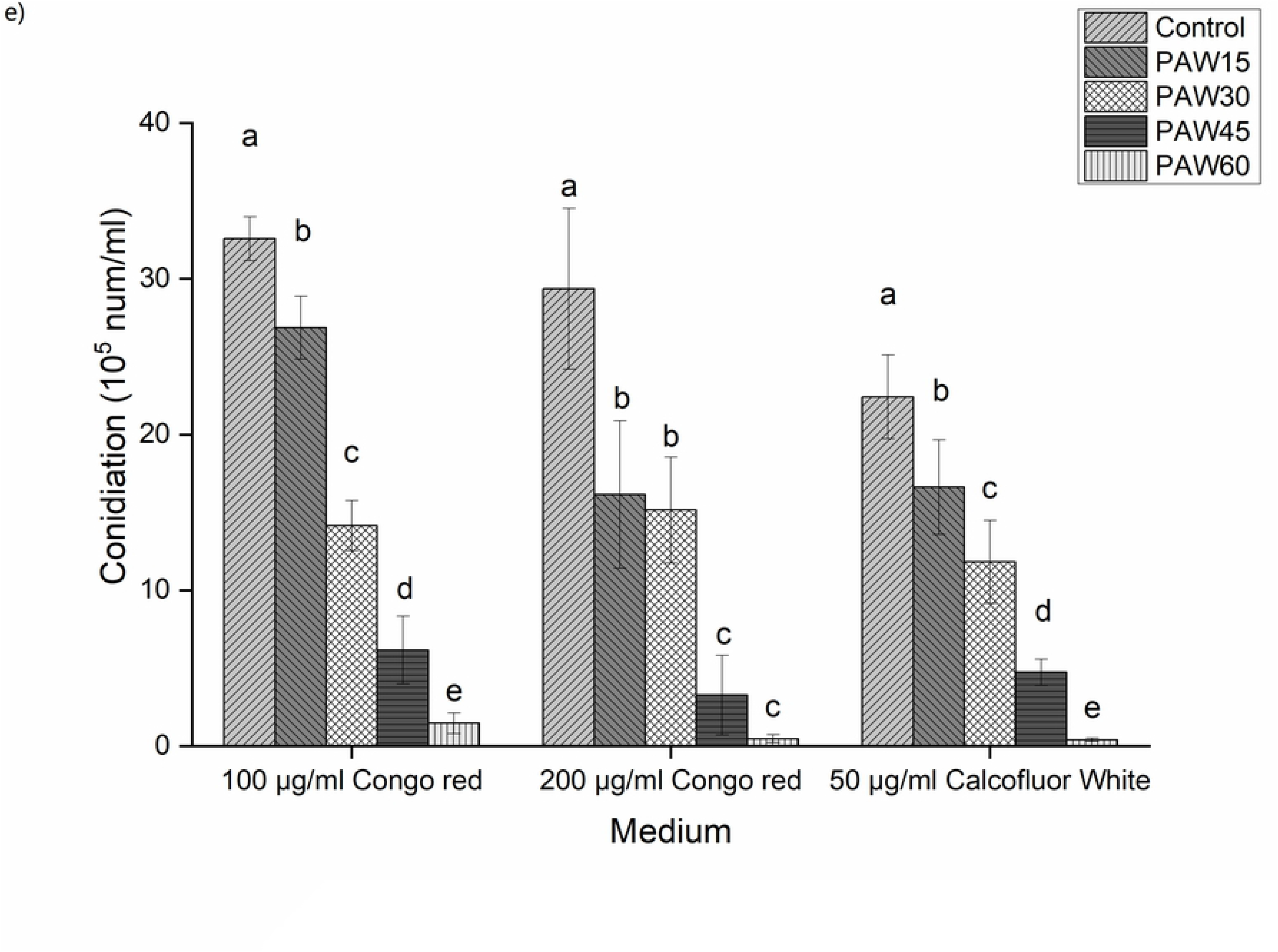
a) Representative image of the colony morphology of *F. graminearum* exposed to PAW15, PAW30, PAW45, and PAW60 treatment for 1h; an image was obtained every 12 h after treatment with PAW during 5 days of incubation at 28 °C. b) Dry mycelial weight after PAW treatment cultured in CM supplemented with 100 mg L^-1^, 200 mg L^-1^ Congo red and 50 mg L^-1^ Calcofluor White. c) Inhibition ratio of mycelium growth after treatment with PAW during 5 days of incubation at 28 °C on CM plate. d) Conidium germination ratio and relative conidium germination ratio of *F. graminearum* after treatment with PAW cultured in CM. R: conidium germination ratio, R_e_: relative conidium germination ratio. e) Conidiation of *F. graminearum* treated with PAW after 7 days culture in CM supplemented with 100 mg L^-1^, 200 mg L^-1^ Congo red and 50 mg L^-1^ Calcofluor White. The results represent the mean±standard deviation (n=6). Vertical bars represent standard deviation of the mean, columns with different letters represent statistically significant results (*P* < 0.05).

PAW treatment could inhibit the germination of *F. graminearum* spores. The pH value of PAW decreased over plasma activation time. However, previous studies demonstrated that pH has no effect on *F. graminearum* germination (Beyer et al., 2004; Depasquale and Montville, 1990). Studies have distinguished PAW-mediated inhibition from inhibition caused by pH alone, and the results have suggested that the inhibitory effect on conidium germination was mainly due to the introduction of PAW. The relative conidium germination ratios of *F. graminearum* treated with PAW15, PAW30, PAW45, and PAW60 are shown in Fig. 3d. Similar results were reported after spores were treated with farnesol, i.e., after 6 h, 44% of them failed to germinate (Semighini et al., 2008). The results of our study also indicated that the antifungal activity of PAW increased over plasma activation time. The conidium germination ratios of *F. graminearum* treated with PAW60 and PAW45 were significantly different from that of control (*P*<0.01). Conversely, the conidium germination ratios of *F. graminearum* treated with PAW30 and PAW15 were not significantly different from the control. The results were consistent with the mycelium growth and conidiation assay findings. The inhibitory effect of PAW was time-specific and increased over activation time.

Conidiation was measured after 7 days of culture on CM supplemented with 100 mg L^-1^or 200 mg L^-1^ Congo red and 50 mg L^-1^ Calcofluor White. As shown in Fig. 3e, the conidial production of PAW treated strains was significantly (*P* < 0.05) reduced compared to that of the nontreated strains after 7 days of incubation. All the groups exhibited suppression of conidiation. The inhibitory effect increased over the plasma activation time. PAW60 had the greatest inhibitory effect on the conidiation of *F. graminearum*, and PAW15 had a weaker inhibitory effect on the conidiation than the other PAW treatment. The results demonstrate that PAW could effectively suppress conidiation.

The total microbial activity of the spores was estimated through the fluorescein diacetate (FDA) method. As shown in Fig. 4c, PAW-treated spores exhibited a decreased proportion of green fluorescence, suggesting a decrease in the total microbial activity of spores after PAW treatments. This result was in line with *in vitro* fungicidal activity assay.

**Fig. 4.**
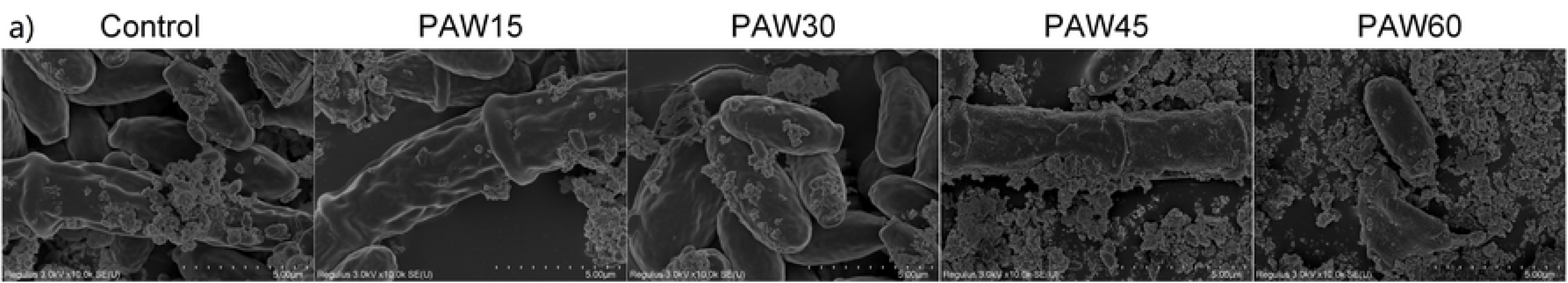
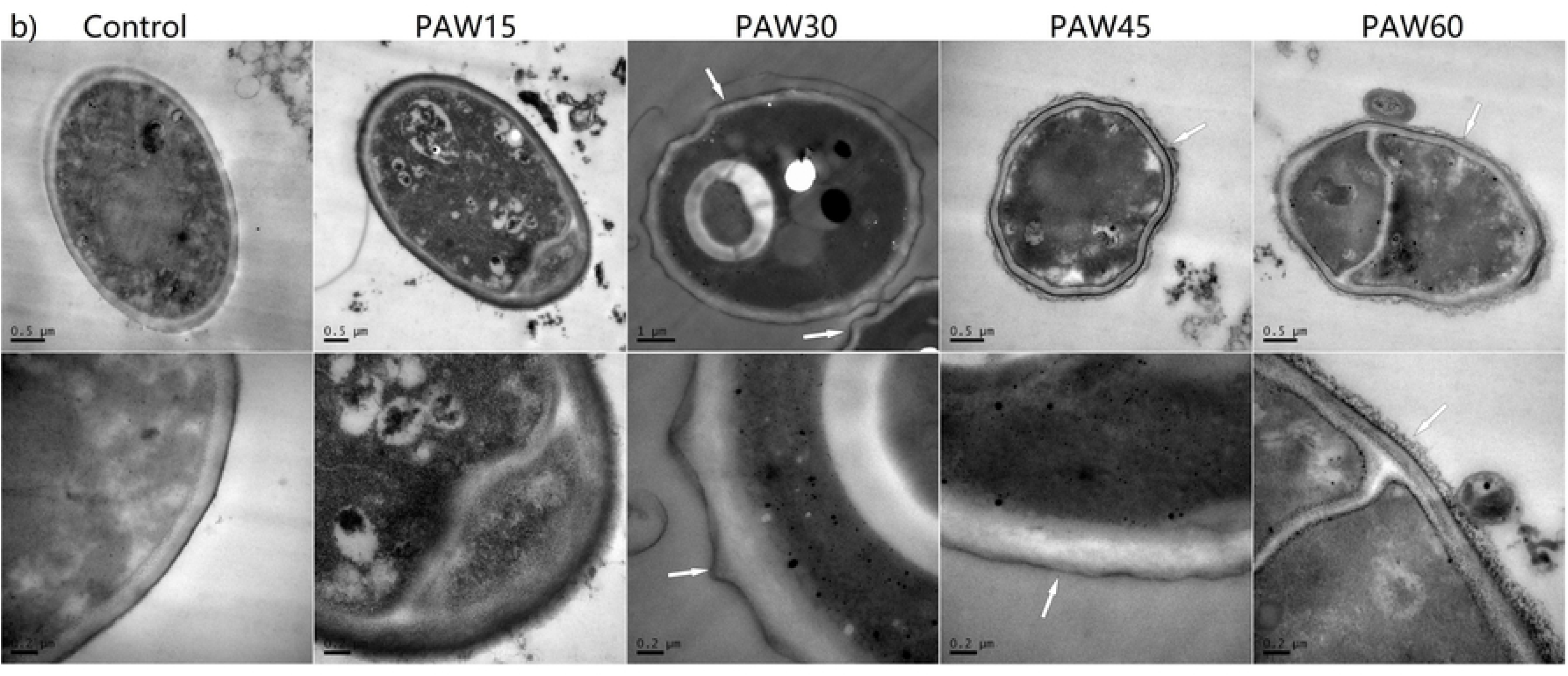
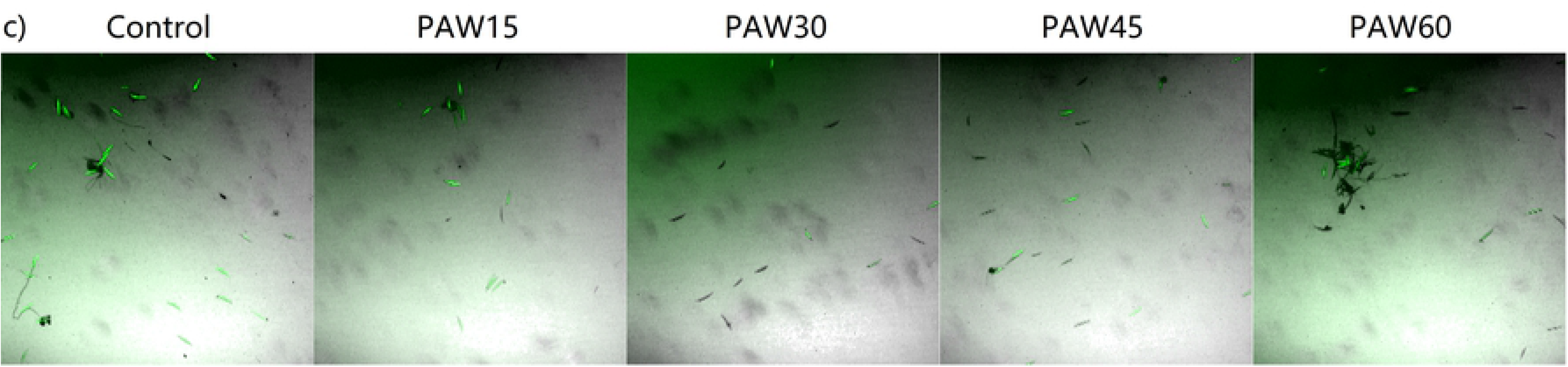
a) SEM images of spores treated with PAW. b) TEM images of spores treated with PAW. c) The fluorescence images of *F. graminearum* spores stained by FDA.

The mode of PAW action in fungi is still subjected to thorough studies, but it has been suggested that biological reactive substances such as ROS and RNS in PAW or cold plasma act synergistically in microbial inactivation (Ercan et al., 2013; Los et al., 2020; Ma et al., 2015; Puligundla et al., 2018; Šimončicová et al., 2018). The fungal cell wall and plasma membrane are the first cell structures that collide with the radicals ROS and RNS in PAW. These reactive species could induce the oxidation of glucose, (N-acetyl)-glucosamine, glycoproteins and glucan in the cell wall and lipid peroxidation in the plasma membrane, causing membrane permeability changes, membrane damage, surface sculpturing in cell wall, and eventually the direct exposure of intracellular components to reactive species. The SEM and TEM results showed that spore cell wall and membrane sculpturing occurred after PAW treatment (Fig. 4), supporting this hypothesis.

In addition to reactive species, the low pH can also inhibit hyphal growth (Wiebe et al., 1996). The intracellular pH value plays an important role in the maintenance of normal cell function, and PAW can cause significant acidification of the intracellular environment. Disruption of intracellular pH homeostasis leads to irreversible cell damage (Lagadic-Gossmann et al., 2004; Xu et al., 2020). The results indicated that acidification of the intracellular environment is mainly attributed to cell wall damage and cell membrane dysfunction. A large amount of protons in PAW can directly enter *F. graminearum* cells through damaged cell membranes.

### 3.4. Effect of PAW on cell morphology and cell membrane integrity

Morphological changes were observed in spores after PAW treatment (Fig. 4a and Fig. 4b). The cell walls were intact in the control spores. No obvious morphological changes were present in the spores treated with PAW15. Morphological changes were observed in the spores treated with PAW30, PAW45 and PAW60. The results from SEM and TEM observations in this study showed that PAW could cause the surface sculpturing in the cell walls of spores, which may be attributed to the oxidative damage induced by reactive species in PAW (Puligundla et al., 2018; Timoshkin et al., 2012; Xu et al., 2020). It is generally agreed that ROS can lead to cell wall damage (Lu, et al., 2017; Shen et al., 2016; Xiang et al., 2018; Xu et al., 2020). The composition of the cell wall, which includes glucose, (N-acetyl)-glucosamine, glycoproteins and glucan, can be oxidized by these reactive species.

The cell membrane is an important organelle that maintains normal cell physiological metabolism. In addition to cell wall damage, reactive species can cause changes in membrane permeability (Gaunt et al., 2006). In this study, the FDA staining method and the DNA/RNA and protein leakage method were employed to assess the effect of PAW on cell membrane integrity (Xiang et al., 2019a; Xu et al., 2020). FDA easily entered the cell membranes and was cleaved by the enzymatic activity of nonspecific esterase and, hence, was detected within live cells, and its fluorescence depends on cell membrane integrity (Grimm et al., 2013; Jones et al., 2016). The decreased proportion of green fluorescence indicated a defect in cell membrane integrity (Fig. 4c).

The leakage of intracellular DNA/RNA and proteins was also a significant indicator of the disruption of cell membrane integrity. As shown in Table 1, upon the addition of PAW to *F. graminearum* spores, there were no significant differences in absorbance (260 nm and 280 nm) in any of the treatment groups compared with the control. Our results indicated that there was no leakage of DNA, RNA or protein, and the cell membranes of the spores were not severely compromised by PAW.

**Table 1.**
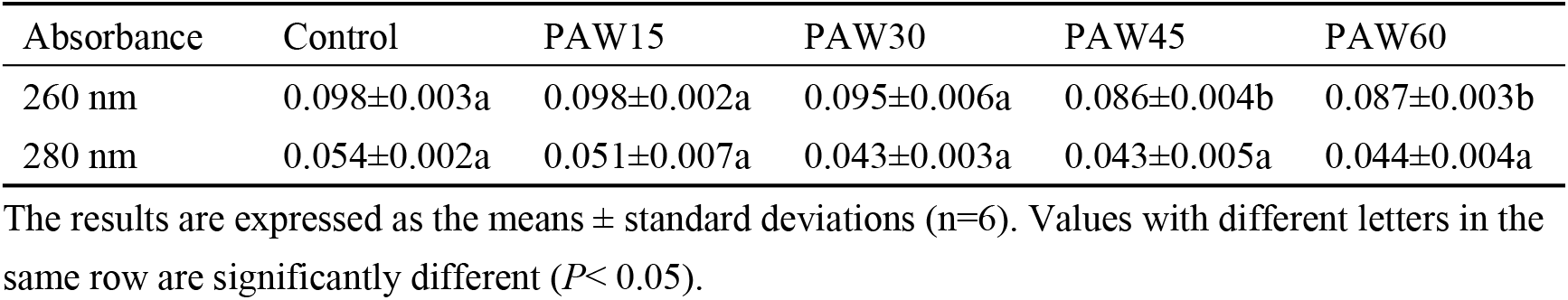
Release of intracellular nucleic acids and protein from spores treated with PAW.

**Table 2.**
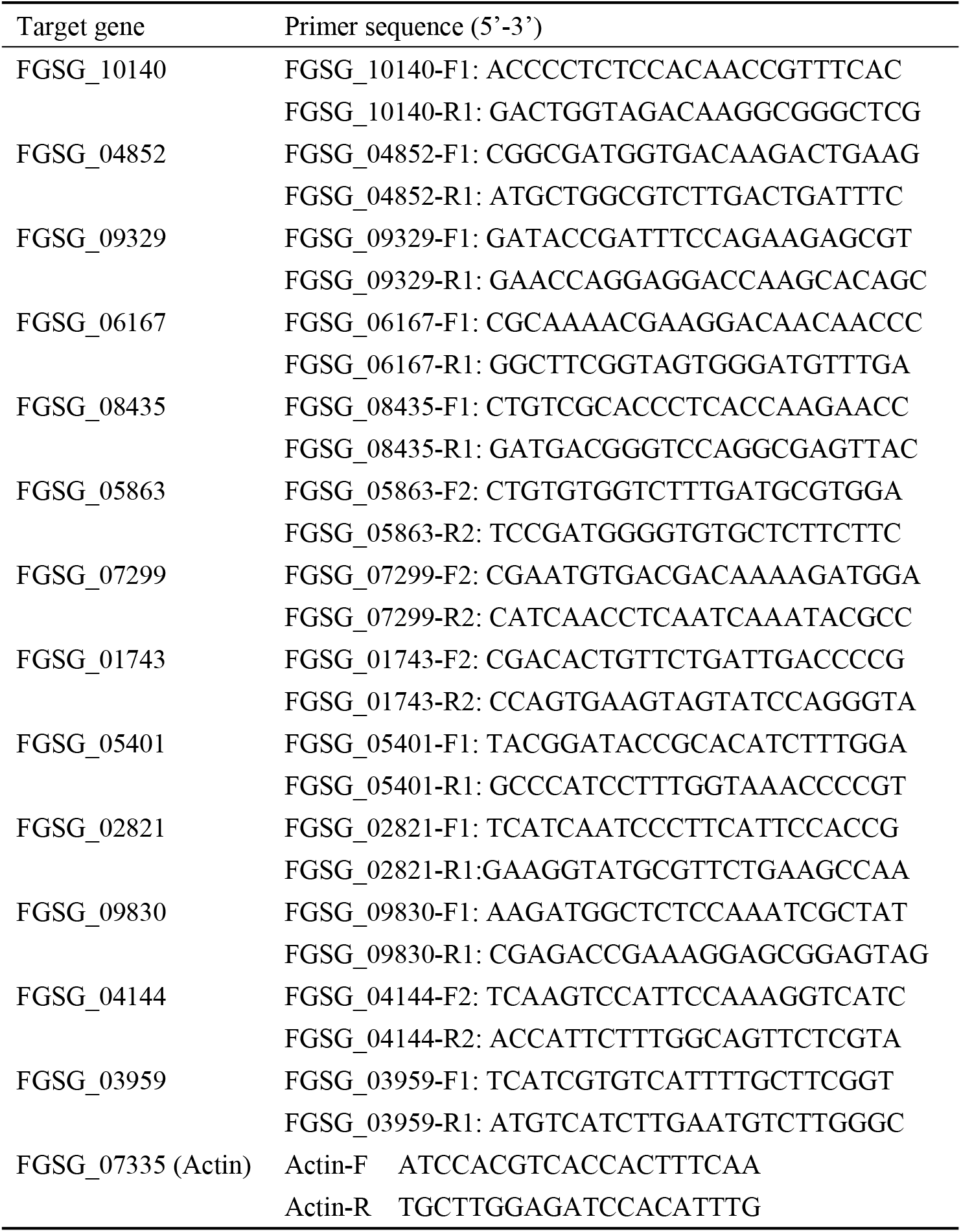
A list of primers used in real-time PCR analysis

In previous literature (Xiang et al., 2018; Xiang et al., 2019a), PAW treatment resulted in the leakage of nucleic acids and proteins in *Pseudomonas deceptionensis* CM2 and *Escherichia coli* O157:H7, which was not identical to our study, the leakage of intracellular DNA/RNA and proteins was not observed in this work. The main reasons for this difference are probably attributed to the difference in membrane composition and the different ORP value of PAW. PAW can change the membrane permeability and allow small inorganic molecules such as RONS to enter the spore, but the extent of the change in membrane permeability was not sufficient to induce the leakage of biomacromolecules in this study.

### 3.5. Measurement of mitochondrial activity

The mitochondrial membrane potential (Δψ_m_) is an indicator of mitochondrial membrane integrity, and Δψ_m_ depolarizes when the membrane is perturbed, consequently combining with less TMRM probe. As shown in

Fig. 5 and Fig. S1**Error! Reference source not found.**, the fluorescence intensity of spores decreased significantly after PAW treatment, indicating that PAW could induce mitochondrial dysfunction, which is an important factor for *F. graminearum* inactivation by PAW. Δѱ_m_ is essential for the survival of the cell as it drives the synthesis of ATP and maintains oxidative phosphorylation (Ly et al., 2003). Furthermore, the degree of depolarization of Δѱ_m_ was in line with the ORP of PAW, and analysis of intracellular redox pairs such as NADH/NAD+ and GSH/GSSG indicated a correlation between extracellular ORP and intracellular redox homeostasis (Li et al., 2019). The existing literatures suggest that a significant increase of intracellular ROS could be induced by PAW (Ma et al., 2013; Tian et al., 2015; Xu et al., 2020). Based on these results, it is hypothesized that extracellular ORP leads to the accumulation of intracellular ROS, and ROS induce the opening of the mitochondrial permeability transition pore (mPTP), causing the depolarization of Δѱ_m_ and mitochondrial dysfunction.

**Fig. 5.**
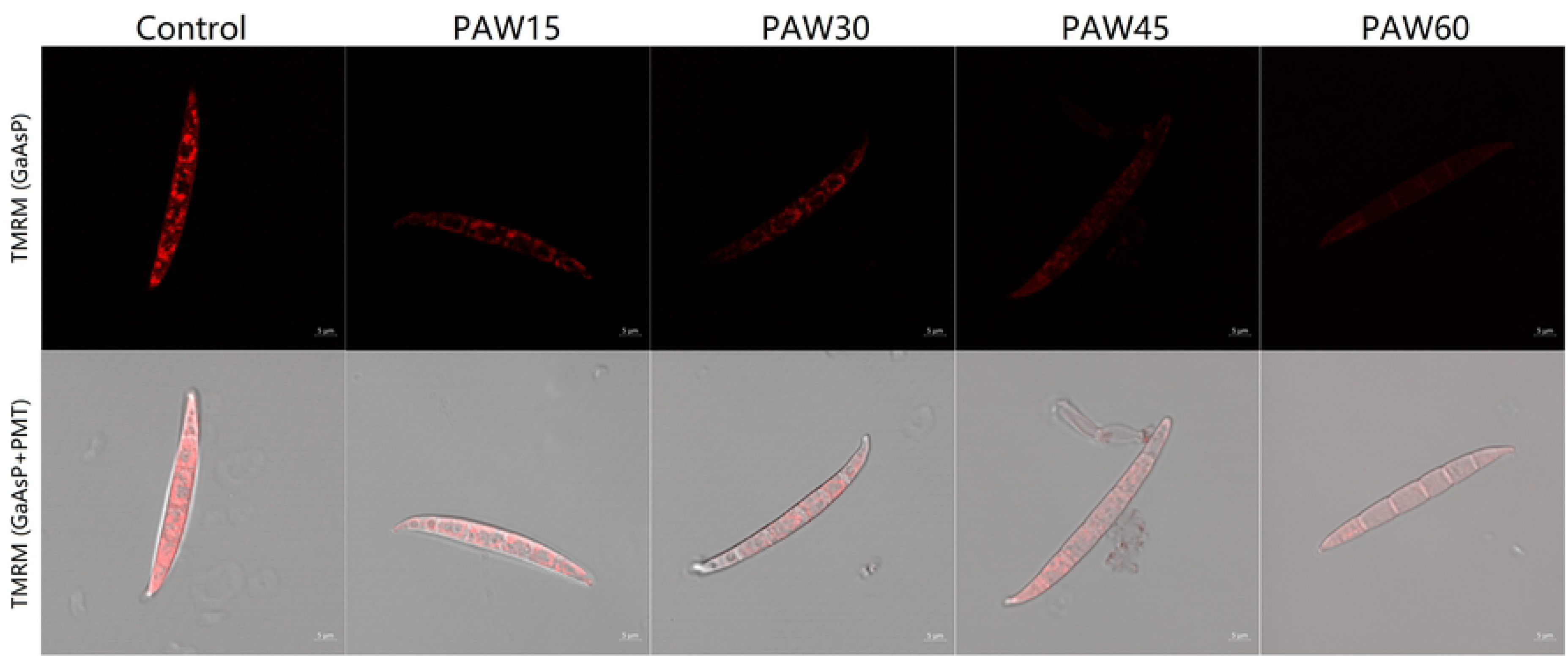
The fluorescence images of *F. graminearum* spores stained by TMRM.

### 3.6. Overview of changes in gene expression of F. graminearum in response to PAW treatment

The numbers of clean reads for PAW-treated and untreated samples were 42,916,050±830015 and 39,063,536±918,515, respectively. The indicator Q30 value for PAW-treated and untreated samples were 94.15±0.15% and 94.37±0.07%, respectively. Uniquely mapped reads in the PAW-treated and -untreated libraries matched 82.49±1.88% and 87.20±1.76% of the total reads, respectively.

### 3.7. Differential gene expression analysis of F. graminearum strain PH-1 treated with PAW

Mapping of the raw RNA-seq expression data revealed that a total of 2837 (22%) of the 12,792 *F. graminearum* unigenes were differentially regulated by PAW treatment. The thresholds for differential expression were *P* adjusted < 0.001 and expression fold Change > 5 for up- and downregulated genes. By these criteria, we identified 703 genes upregulated and 676 genes downregulated by PAW. As shown in Fig. 6, of the 1379 genes, 944 genes were classified as having “unknown function”.

**Fig. 6.**
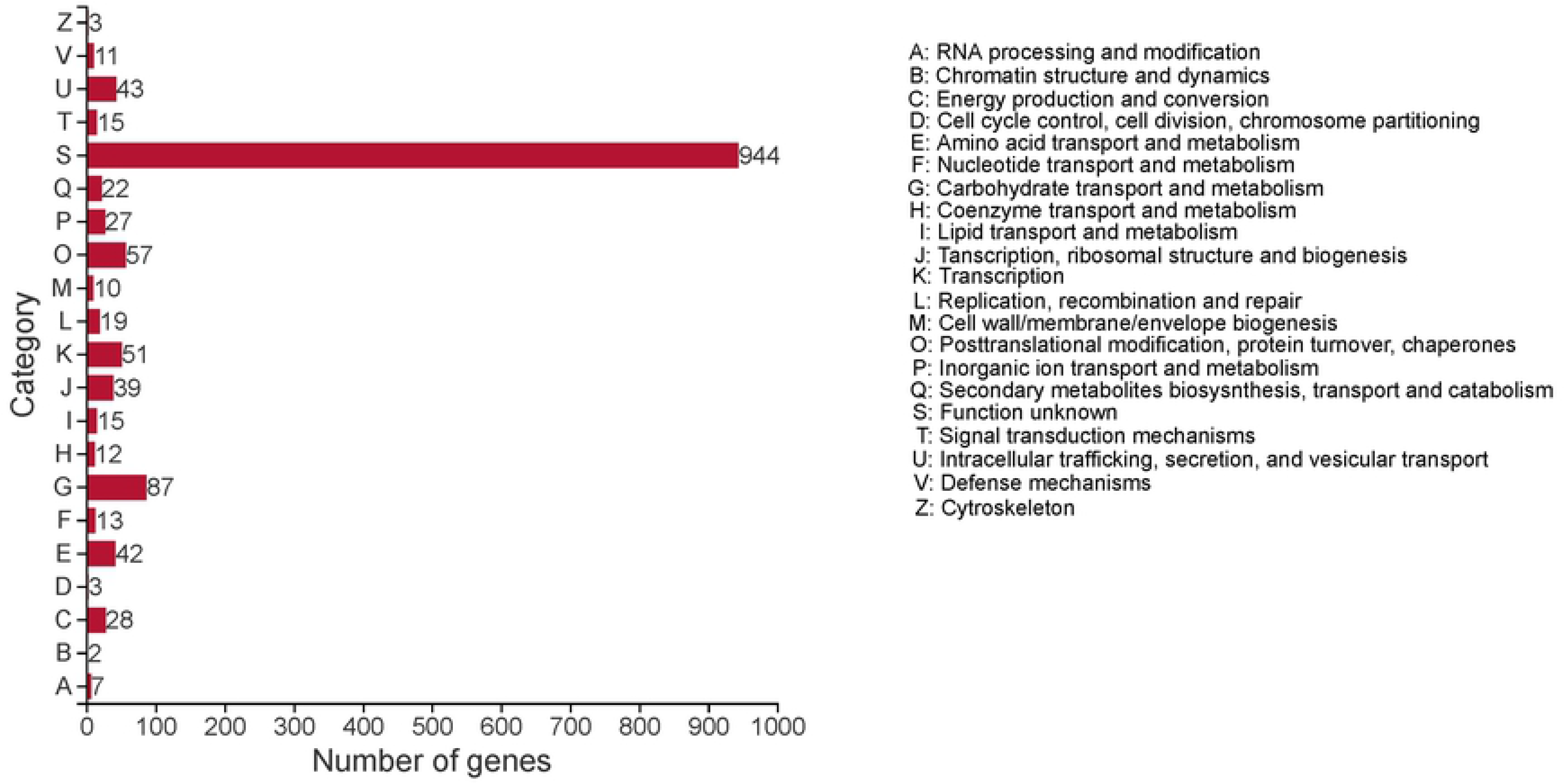
COG categories of differentially expressed genes relative to PAW-treated *F. graminearum*.

There were 3 putative genes (FGSG_10140, FGSG_04852, and FGSG_09329) involved in mitochondrial function in the 10 most upregulated genes and 1 gene (FGSG_03959) involved in the cell membrane in the top 5 downregulated genes in response to PAW. The results indicated that the cell membrane and mitochondria were the organelles most affected by PAW. Based on the results of this report and previous literatures, we mainly focused on the genes involved in mitochondrial function and cell wall and membrane integrity (Liu et al., 2010; Suwal et al., 2019; Los et al., 2020; Xu et al., 2020).

The most upregulated gene involved in mitochondrial function is listed in Table 3. The differences in gene expression of treated PH-1 ranged from up to a 3662.08-fold increase to a 11.35-fold increase compared to untreated PH-1. The most upregulated gene was FGSG_10140, and the putative product encoded by the gene is succinate dehydrogenase assembly factor. It is well known that succinate dehydrogenase (complex II; or succinate: ubiquinone oxidoreductase, SQR) is a functional member of the aerobic respiratory chain. Complex II couples the oxidation of succinate to fumarate in the mitochondrial matrix with the reduction of ubiquinone in the membrane (Yankovskaya et al., 2003). It was found that the SQR redox centers are arranged in a manner that aids the prevention of ROS formation at the flavin adenine dinucleotide. This is likely to be the main reason FGSG_10140 and other genes are highly expressed during PAW treatment, which can induce high intracellular ROS levels.

**Table 3.** Response to PAW of the genes involved in mitochondrial function

| Accession number | Putative product encoded by the gene | Fold change in gene expression<sup>a</sup> | Padjust |
| --- | --- | --- | --- |
| FGSG_10140 | Hypothetical protein similar to succinate dehydrogenase assembly factor 2, mitochondrial | 3662.08 | <0.001 |
| FGSG_04852 | Hypothetical protein similar to NADH-cytochrome b5 reductase | 2195.27 | <0.001 |
| FGSG_09329 | Hypothetical protein similar to ATPase | 2122.66 | <0.001 |
| FGSG_06167 | Hypothetical protein similar to 54s ribosomal protein L4, mitochondrial | 839.12 | <0.001 |
| FGSG_08435 | Hypothetical protein similar to nitronate monooxygenase | 783.80 | <0.001 |
| FGSG_05863 | Mitochondrial import protein 1 | 714.07 | <0.001 |
| FGSG_07299 | Hypothetical protein similar to transferase CAF17, mitochondrial | 37.54 | <0.001 |
| FGSG_01743 | Acetyl-coenzyme A synthetase | 11.35 | <0.001 |

<sup>a</sup>Fold-change value represents the fold expression in *F. graminearum* treated with PAW as compared to that in nontreatment control

The top up- or downregulated genes involved in cell wall or membrane integrity are listed in Table 4. The differences in gene expression of treated PH-1 ranged from up to a 1731.37-fold increase to a 0.00049-fold decrease compared to untreated PH-1. The most upregulated gene was FGSG_02821, and the putative product encoded by the gene is a transmembrane protein that belongs to the HPP family according to NCBI BLAST analysis. These proteins are integral membrane proteins. While the function of these proteins is uncertain, they may be transporters. It has been shown that the HPP family of integral membrane proteins transports nitrite across the chloroplast membrane (Krapp, 2015; Maeda et al., 2014). In addition, nitrate and nitrite exist in PAW generated by air plasma. Based on these results, it is hypothesized that the FGSG_02821 gene is upregulated mainly in response to high nitrite or nitrate levels in PAW.

**Table 4.** Response to PAW of the genes involved in cell wall and membrane integrity

| Accession number | Putative product encoded by the gene | Fold change in gene expression<sup>a</sup> | Padjust |
| --- | --- | --- | --- |
| FGSG_02821 | Hypothetical protein similar to transmembrane protein | 1731.37 | <0.001 |
| FGSG_04144 | Hypothetical protein similar to vi polysaccharide biosynthesis vipa tvib | 47.51 | <0.01 |
| FGSG_05401 | Hypothetical protein similar to Beta-1,3-glucanase | 12.46 | <0.001 |
| FGSG_09830 | C-4 methylsterol oxidase | 9.60 | <0.001 |
| FGSG_03959 | Hypothetical protein similar to plasma membrane protein pth11 | 0.00049 | <0.001 |

<sup>a</sup>Fold-change value represents the fold expression in *F. graminearum* treated with PAW as compared to that in nontreatment control

The most downregulated gene was FGSG_03959, and the putative product encoded by the gene was protein PTH11 according to NCBI BLAST analysis. PTH11 has been shown to be important for appressorium formation and pathogenicity in *Magnaporthe grisea* (DeZwaan et al. 1999; Kou et al., 2016). In addition, 30 genes were detected during infection of three hosts (wheat, barley, and maize) encoding G-protein coupled receptors (GPCRs) belonging to the integral membrane protein PTH11 class, and many exhibited host-preferential expression (Harris et al., 2016). As shown in Supplementary Table 1, 11 of the 30 genes showed differential expression during PAW treatment in our work, and 7 of the 11 genes were downregulated genes. Deletion of these downregulated genes resulted in a significant reduction in conidiation, confirming the result that PAW treatment reduces the conidiation of *F. graminearum*. However, the most downregulated gene, FGSG_03959, showed no or barely detectable expression in hyphae, perithecia and infected wheat heads during infection, indicating that the gene is probably not essential for plant infection (Jiang et al., 2019).

### 3.8. Validation of the differential expression results

In line with our main objective of identifying cell wall-, membrane integrity- or mitochondria-related genes, thirteen PH-1 unigenes were selected for further analysis on the basis of their expression levels and possible roles in defense mechanisms in response to PAW (Table 2). As shown in Table 5, in general, the Q-PCR results correlated with the transcriptomic data. However, the fold changes in expression determined by DESeq2 sequencing were not in line with those determined by Q-PCR. We observed that the expression levels of FGSG_10140, FGSG_04852, FGSG_09329, FGSG_06167, FGSG_08435, FGSG_05863 and FGSG_02821 determined by DESeq2 analysis were much higher than those determined by Q-PCR. The possible reason for these discrepancies is that the precision and accuracy of the methods are different.

**Table 5.** Comparison in the changes of gene expression determined by DESeq2 sequencing and real-time PCR approaches

| Accession number | Putative product encoded by the gene | Fold change in gene expression determined by DESeq2 sequencing<sup>a</sup> | Fold change in gene expression determined by real-time PCR<sup>a</sup> |
| --- | --- | --- | --- |
| FGSG_10140 | Hypothetical protein similar to succinate dehydrogenase assembly factor 2, mitochondrial | 3662.08 | 52.83 |
| FGSG_04852 | Hypothetical protein similar to NADH-cytochrome b5 reductase | 2195.27 | 40.41 |
| FGSG_09329 | Hypothetical protein similar to ATPase | 2122.66 | 2.52 |
| FGSG_06167 | Hypothetical protein similar to 54s ribosomal protein L4, mitochondrial | 839.12 | 14.32 |
| FGSG_08435 | Hypothetical protein similar to nitronate monooxygenase | 783.80 | 22.12 |
| FGSG_05863 | Mitochondrial import protein 1 | 714.07 | 12.64 |
| FGSG_07299 | Hypothetical protein similar to transferase CAF17, mitochondrial | 37.54 | 2.18 |
| FGSG_01743 | Acetyl-coenzyme A synthetase | 11.35 | 19.47 |
| FGSG_02821 | Hypothetical protein similar to transmembrane protein | 1731.37 | 7.95 |
| FGSG_04144 | Hypothetical protein similar to vi polysaccharide biosynthesis vipa tvib | 47.51 | 7.13 |
| FGSG_05401 | Hypothetical protein similar to Beta-1,3-glucanase | 12.46 | 10.90 |
| FGSG_09830 | C-4 methylsterol oxidase | 9.60 | 13.48 |
| FGSG_03959 | Hypothetical protein similar to plasma membrane protein pth11 | 0.00049 | 0.080 |

<sup>a</sup>Fold-change value represents the fold expression in *F. graminearum* treated with PAW as compared to that in nontreatment control

## 4. Conclusion

In conclusion, the results from our study confirmed that PAW treatment is a highly effective disinfection procedure against *F. graminearum*. It has the potential to control fungal contamination. This study also unravels the potential antifungal mechanism of PAW from the perspective of cellular response and differential gene expression. ROS and RNS in PAW first compromised the cell membrane, leading to intracellular ROS accumulation, and then intracellular pH decreased, and depolarization of Δѱ_m_ and mitochondria dysfunction occurred. The DESeq2 sequencing analysis enhanced the hypothesis by the fact that there were three putative genes involved in mitochondrial function in the 10 most upregulated genes and one gene involved in the cell membrane in the top 5 downregulated genes in response to PAW treatment. The information obtained from this work may verify the feasibility and validity of the application of this technique in plant disease control and provides important insights into the antifungal mechanism of PAW to fight against *F. graminearum*.

## Acknowledgment

The authors gratefully acknowledge the financial support provided by the National Natural Science Foundation of China [grant number 31900126 to L. Li and 31701723 to J. Guo]; Zhejiang key research and development program [grant number 2018C02G2011099]; and Program of Innovative Entrepreneurship Training for Undergraduate of Zhejiang A & F University [grant number 2020KX0002, 2020KX0025].

